# Circularization of *rv0678* for genotypic bedaquiline resistance testing of *Mycobacterium tuberculosis*

**DOI:** 10.1101/2022.10.05.511060

**Authors:** Jason D Limberis, Alina Nalyvayko, Joel D Ernst, John Z Metcalfe

## Abstract

Circular DNA offers benefits over linear DNA in diagnostic and field assays, but currently, circular DNA generation is lengthy, inefficient, highly dependent on the length and sequence of DNA, and can result in unwanted chimeras. We present streamlined methods for generating PCR-targeted circular DNA from a 700bp amplicon of *rv0678*, the high GC content (65%) gene implicated in *Mycobacterium tuberculosis* bedaquiline resistance, and demonstrate that these methods work as desired. We employ self-circularization with and without splints, a Gibson cloning-based approach, and novel two novel methods for generating pseudocircular DNA. The circular DNA can be used as a template for rolling circle PCR followed by long-read sequencing, allowing for the error correction of sequence data, and improving the confidence in the drug resistance determination and strain identification; and ultimately improving patient treatment.

## Introduction

Circular DNA offers many benefits over linear DNA in diagnostic and field assays. The advantages primarily derive from circular DNAs’ resistance to most exonucleases – which catalyze the removal of nucleotides from the free ends of single-, or double-stranded DNA by hydrolyzing phosphodiester bonds, and its ability to act as a template for rolling circle amplification – the isothermal process of unidirectional nucleic acid replication resulting in concatenated copies of the circular template. Rolling circle amplification utilized in assays detecting vascular disease-related SNPs in clinical samples^1^, ultrasensitive isothermal protein analyte detection in microscale platforms^2^, in-situ RNA analyte detection^3^, and magnetic and optomagnetic sensor systems^4–6^, but remains unexplored for molecular tuberculosis tests.

Rolling circle amplification was first developed as a method in the mid-1990s^7–11^. A typical application of rolling circle amplification is generating long, single-stranded DNA concatemers as the template for long-read sequencing as with Oxford Nanopore Technologies (ONT) platforms, allowing error correction by taking a consensus of the de-concatenated sequence^12^. Sequencing of longer DNA strands also improves the output from the sequencer, as short reads exhaust nanopores more quickly. ONT sequencing is highly sought after as this technology is portable and requires little infrastructure, allowing for its utilization in the field, for example, in the characterization of unique ecological niches^13^. It also has a potentially pioneering impact on diagnostics and clinical practice, for example, in the point-of-care genotypic drug susceptibility testing of pathogens, such as *Mycobacterium tuberculosis*^14^, for which no comprehensive point-of-care drug resistance test exists.

Current methods of generating circular DNA are lengthy, inefficient, highly dependent on the length and sequence of DNA, and can result in unwanted chimeras^15,16^. The circularization of single-stranded DNA is the most common^17–24^, and is used to generate templates for short-read sequencing in BGI’s nanoball technology^25^. A second approach is to clone a fragment into a vector, generating circular DNA; however, this will always result in the presence of the vector backbone. A third approach is to ligate dumbbell (hairpin) oligos to linear dsDNA as utilized by PacBio in their Single Molecule, Real-Time (SMRT) sequencing system. The pseudo-circularized DNA then serves as a rolling circle amplification template.

Here, we present streamlined methods for generating PCR (or similar techniques, including strand displacement and recombinase polymerase amplification) targeted circular DNA. We focus on generating a ~700bp amplicon of *rv0678*, the high GC content (65%) gene implicated in bedaquiline resistance in *M tuberculosis*, the causative agent of tuberculosis. We demonstrate several methods, including using splints, a Gibson cloning-based approach for self-circularization, and novel methods for generating pseudo-circular DNA.

## Methods

### Amplicon generation

We generated the initial ~700bp amplicon using the primer set “initial amplicon generation” (**Table 1**) from genomic *M. tuberculosis* H37Rv DNA with Q5 polymerase (NEB, USA) according to the manufacturer’s instructions. The thermocycling was done as follows: initial denaturation at 98°C for 30 seconds, 34 cycles of 98°C, 62°C, and 72°C for 10, 10, and 20 seconds respectively. Amplicons were purified using 0.8X Agencourt AMPureXP beads (BD, USA) according to the manufacturer’s instructions. This template was then used, with the primers in **Table 1** to generate the remaining amplicons using the same procedure.

**Table 1:**
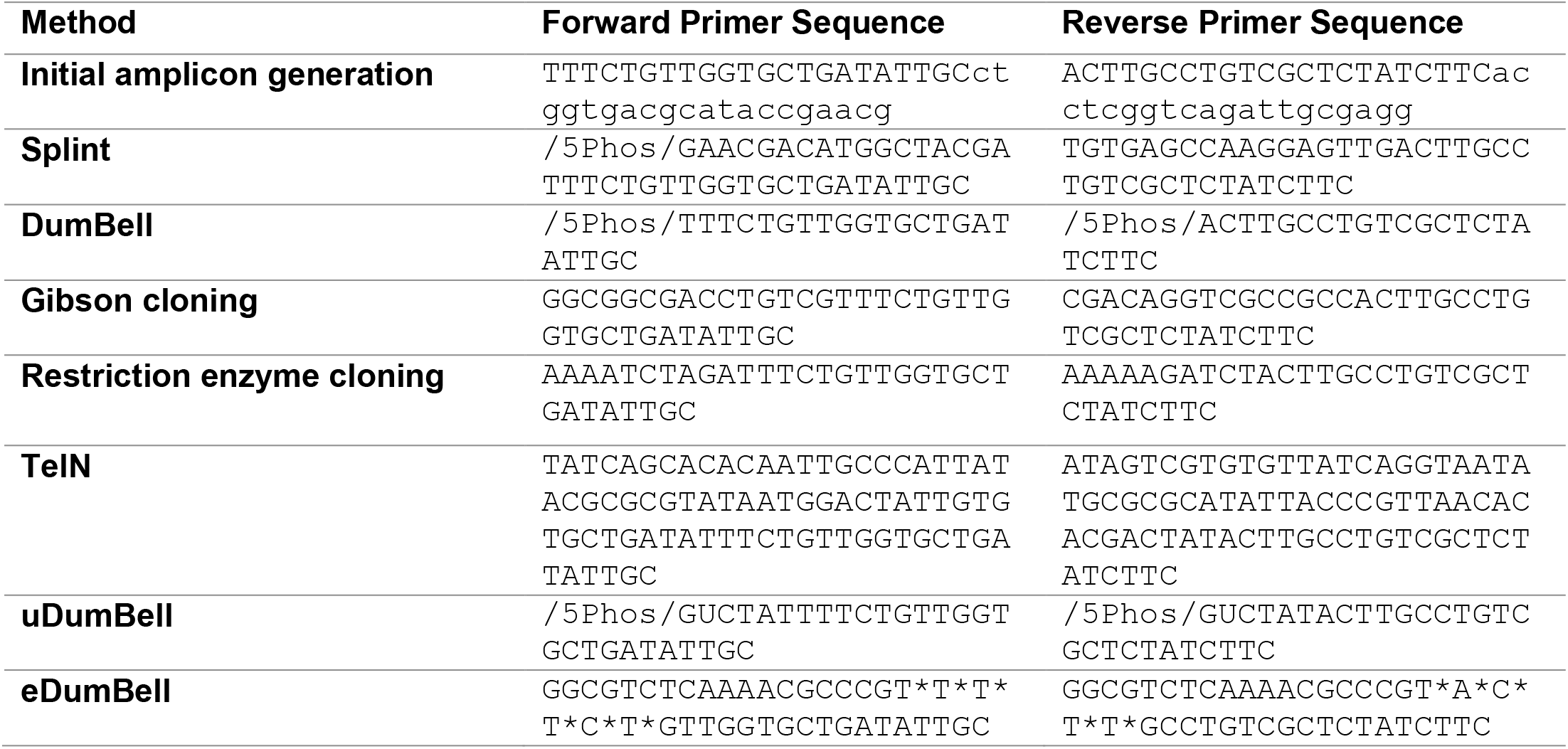
Primer sequences used in this study. An asterisk between nucleotides denotes a phosphorothioate bond.

### DNA circularization procedures

#### Splint

Two micrograms of amplicon was digested with Lambda Exonuclease (NEB, USA) in a 30ul reaction at 37°C for 30 minutes according to the manufacturer’s instructions. The reaction was stopped by adding EDTA to 20 mM and incubating at 75°C for 10 minutes. Following a 1.8X AMPureXP bead cleanup, T4 Polynucleotide Kinase (NEB, USA) was used to phosphorylate the 5’-end according to the manufacturer’s instructions. A 1.8X AMPureXP bead cleanup was done, and 100ng of the resulting single-stranded material was used to generate single-stranded circular DNA described as follows.

CircleLigase. One hundred units of CircleLigase was used in a 20ul reaction set up according to the manufacturer’s instructions. The reaction was incubated at 60°C for four hours, followed by the inactivation of the enzyme at 80°C for 10 minutes.

Splint ligation. Two nanomolar of the splint (ACACTCGGTTCCTCAACGAACGACATGGCTACGA; **Supplementary Figure 3**) was incubated with the amplicons in a reaction without the ligase or buffer addition, at 80°C for 5 minutes and slowly cooled to 4°C using a ramp rate of 0.1C/s. Either T4 ligase (NEB, USA), Ampligase (Biosearch Technologies, UK), or DNA ligase (NEB, USA) and the corresponding buffer were added and incubated as follows. For the T4 ligase reaction, 22°C, 15°C, 4°C for 30, 120, and 120 minutes; for the Ampligase reaction, 60°C, 55°C, 45°C for 10, 10, and 120 minutes; and for the Taq DNA ligase reaction, 70°C, 65°C, gradient with ramp rate of 0.1C/s to 60°C for 10, 10, and 90 minutes.

*DumBell and uDumBell^26^ (temporary DOI: https://www.protocols.io/private/4AC47731360F11ED9A700A58A9FEAC02)* The dumbbell (hairpin) adapters (`/5Phos/CGAGACAGTAGAAGACCATGAACAAGCAGCACACGATAAACTAGACACCCTACTGTCTCG` and `/5Phos/ATAGACCGAGACAGTAGAAGACCATGAACAAGCAGCACACGATAAACTAGACACCCTACTGTCTCG`; **Supplementary Figure 1**) were prepared by incubating at 80°C followed by cooling to room temperature over 30 minutes. Two hundred nanograms of the amplicon was incubated with 1um of the adapter with T4 ligase in a 30ul reaction at 22°C, 15°C, 4°C for 30, 120, and 120 minutes and inactivated at 65°C for 5 minutes.

#### Gibson

Four hundred nanograms of the amplicon was incubated with NEBuilder HiFi DNA Assembly Master Mix (NEB, USA) in a 20ul reaction at 50°C for 60 minutes.

#### Restriction Enzyme

Eight hundred nanograms of the amplicon was digested with Xball (NEB, USA) at 37°C for 30 minutes, followed by a 1X AMPureXP bead cleanup. Two hundred nanograms of this was incubated with the amplicons in a reaction without the ligase or buffer addition, at 80°C for 5 minutes and slowly cooled to 4°C using a ramp rate of 0.1C/s. Buffer and either T4 ligase, Ampligase, or Taq DNA ligase were added and incubated as described in *Splint ligation* (**Supplementary Figure 2**).

#### Kinase-Ligase

Four hundred nanograms of the amplicon was incubated with KLD Mix (NEB, USA) in a 20ul reaction at room temperature for 30 minutes.

#### TelN

One hundred nanograms of the amplicon was incubated with 10U of TelN Protelomerase (NEB, USA) according to the manufacturer’s instructions at 30°C for 30 minutes, followed by inactivation at 75°C for 5 minutes (**Supplementary Figure 2**).

*easyDB*^27^ *(temporary DOI: https://www.protocols.io/private/36E3DB05360F11ED9A700A58A9FEAC02)*Four-hundred nanograms of the amplicon was incubated with easyDB buffer containing 0.05U of T7 exonuclease, 0.03U of Phusion polymerase, and 53U Taq DNA ligase, 110mMmM Tris-HCl pH 7.5, 15mM MgCl2, 0.4mM dGTP, 0.1mM dATP, 0.1mM dTTP, 0.1mM dCTP, 4mM DTT, 4% PEG 8000, and 0.2mM NAD+ (all sourced from NEB, USA) in a 20ul reaction at 50°C for 60 minutes.

#### Exonuclease treatment to remove non-circular and non-pseudo circular DNA

Twenty microliter reactions containing 10U Exonuclease VIII truncated (NEB, USA) were set up according to the manufacturer’s instructions and incubated at 37°C for 30 minutes. The reaction was inactivated by adding 24mM EDTA and incubating at 70°C for 30 minutes. A 1.8X AMPureXP bead cleanup was then done. For the easyDB method, 50U of Exonuclease III was also included in the reaction, and the reaction was incubated at 37°C for 1 hour before enzyme inactivation.

#### TapeStation

Samples were run on the Agilent TapeStation (Agilent, USA) using the D1000 kit according to the manufacturer’s instructions.

## Results

We generated ~700nt amplicons spanning the *rv0678* gene from *M. tuberculosis* H37Rv DNA as a template for the various methods using the primer set “initial amplicon generation” in **Table 1**. We then generated various amplicons from this template using the remainder of the primer pairs in **Table 1**. Following the various incubations (**Figure 1**), specific exonucleases were used to eliminate non-circular or non-pseudocircular DNA.

**Figure 1:**
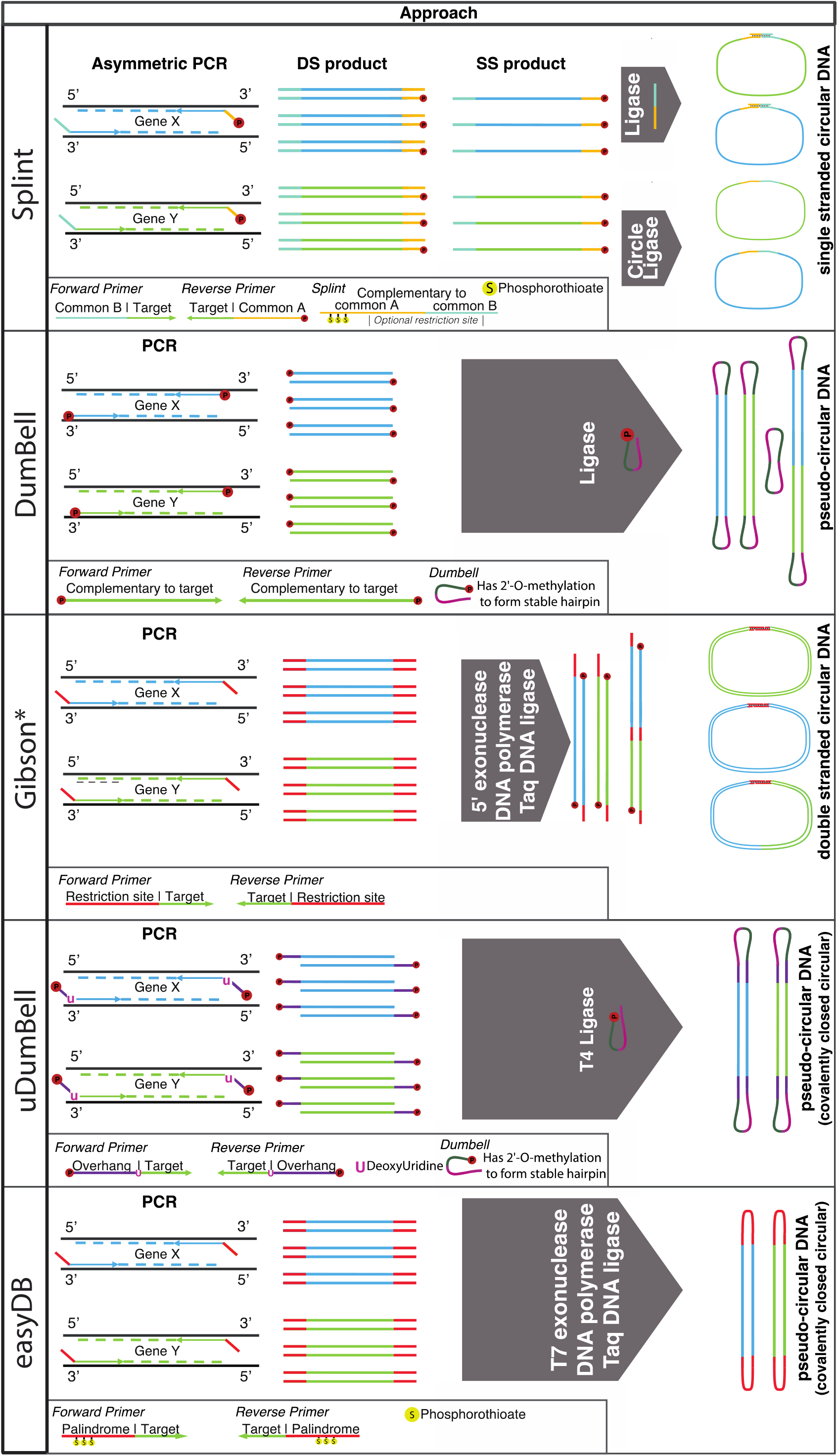
Streamlined methods for the circularization of PCR amplicons.

### Single-stranded circular DNA

Previously described single-stranded DNA circularization methods use a specialized ligase, CircleLigase, an enzyme that is most effective on short (<200nucleoties) strands. Here, we simplified the generation of single-stranded DNA of a large, high GC-content amplicon and attempted self-circularization using CircleLigase and common ligases with an oligonucleotide splint.

Self-circularization of phosphorylated single-stranded DNA using CircleLigase was successful; however, it appears inefficient (**Figure 2** – Splint). Circularization using a splint complementary (**Supplementary Figure 3**) to the two ends of the single-stranded DNA and incubation with either T4 ligase, Ampligase, or Taq DNA ligase similarly resulted in low concentrations of single-stranded circular DNA, with T4 ligase performing the poorest.

**Figure 2:**
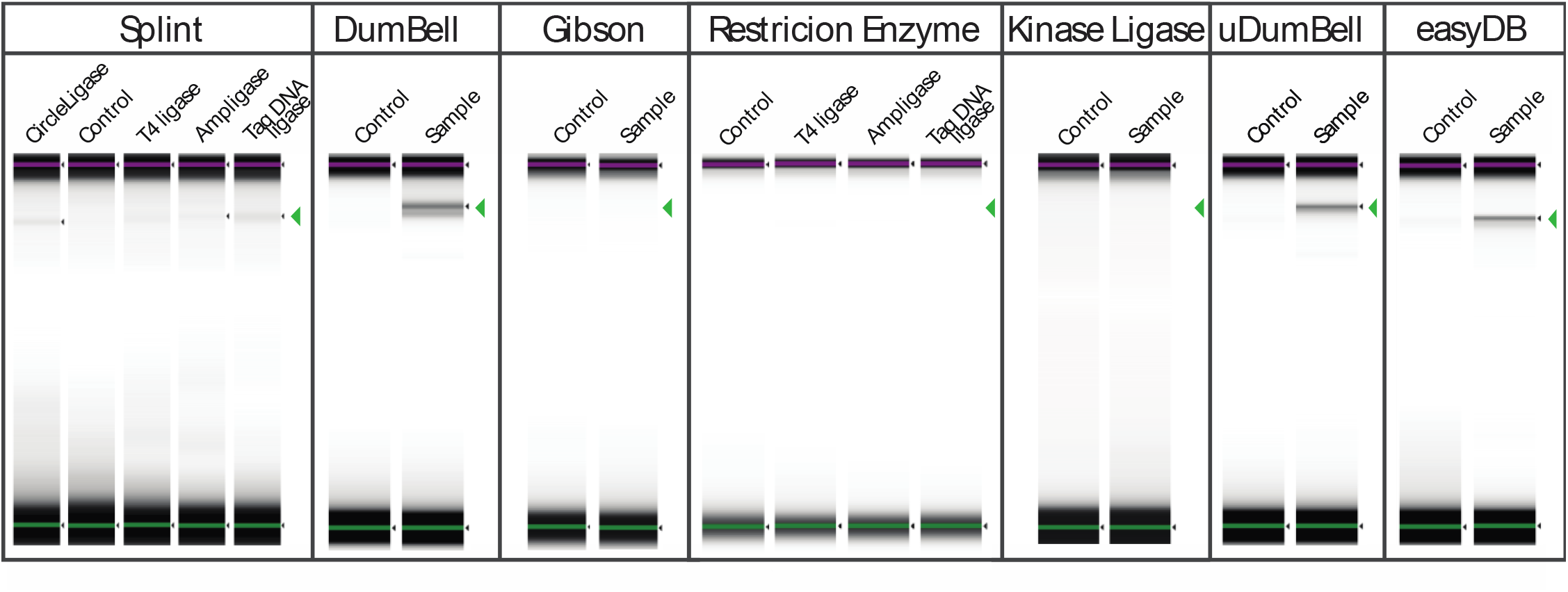
Streamlined methods for circularizing a ~700bp high GC amplicon. Successful circularization was achieved for self-circularization of ssDNA using CircleLiagse, using splints and T4 ligase, Ampligase, and Taq DNA ligase; DumBell ligation using T4 ligase, uDumBell ligation and easyBD ligation. The results panel shows the TapeStation gel image for the control (C) and sample (S) which were treated with specific exonucleases to remove all noncircular or non-pseudocircular DNA; the green arrow indicates the expected band location. We could not generate circularized products using a Gibson cloning-based approach or a restriction enzyme approach using T4 ligase, Ampligase, or Taq DNA ligase. The Kinase ligase approach did not form a single amplicon circular DNA, but we detected large circular concatemers (not visible in this figure).

### Double-stranded circular DNA

Commo methods of double-stranded DNA circularization are used in cloning, where a double-stranded fragment is inserted into a large double-stranded backbone. If the large backbone is not dephosphorylated, it will often self-recircularize. Here we attempted to self-circularize a large, high GC content amplicon.

We generated amplicons with fifteen complementary bases at each end for self-circularization using the Gibson cloning, and kinase, ligase treatment reactions. The Gibson reaction resulted in no detectable circular DNA, while the kinase-ligase treatment did not result in a single circularized amplicon, but it did produce large, concatenated circles. We also generated amplicons with commentary XbalI sites at the ends and digested the amplicons accordingly, which we incubated with T4 ligase, Ampligase, or Taq DNA ligase. In all cases, we did not observe any double-stranded circular DNA. We further attempted using the restriction sites NdeI and KpnI, and a twenty-seven-base pair *insert* with no success.

### Double-stranded pseudo-circular DNA

The ligation of dumbbell (hairpin) oligos to linear dsDNA is used by PacBio to create double-stranded pseudo-circular DNA. We attempted this method, and two streamlined variations to produce double-stranded pseudocircular DNA of a large, high GC content amplicon.

Ligation of DumBells – single-stranded, phosphorylated dumbbell (hairpin) DNA – to double-stranded phosphorylated DNA following DNA purification was successful but did produce concatemerized amplicons and circular DNA formed from only dumbbell (hairpin) DNA (**Figure 2** – DumBell).

Including deoxyUridine in the PCR primer sequences causes Q5 and other high-fidelity polymerases to arrest elongation. This results in overhangs that were successfully ligated to a complementary dumbbell (hairpin) D structure (**Figure 2** – uDumBell). The deoxyUridine reduced the PCR product by approximately two-thirds, but this was ameliorated by increasing the Q5 DNA polymerase concentration three-fold.

We designed primers with a tail sequence that forms a six-nucleotide hairpin at temperature <55°C, but not ≥55 °C (**Supplementary Figure 4**).These primers contain six phosphorothioate bonds starting at the complementary region to inhibit exonuclease T7 activity. The primers successfully amplified the target and, following incubation with a mixture of T7 exonuclease, DNA polymerase, and Taq DNA ligase, pseudo-circular double-stranded DNA formed (**Figure 2** – <u>easyDB</u>).

We also made pseudo-circular DNA using TelN protelomerase, which cuts dsDNA at a 56bp recognition sequence and leaves covalently closed ends at the cleavage site.

## Discussion

Many life forms utilize circular DNA; for example, bacterial chromosomes and plasmids, and the eukaryotic mitochondrial genome are made of circular double-stranded DNA. Numerous viruses use covalently closed circular DNA and rolling circle amplification, including Phi X174, hepadnaviruses (e.g., hepatitis B), herpesvirus, and poxviruses ^28^. The poxviruses have a pseudo-circular genome in which linear double-stranded DNA is covalently closed at the ends forming hairpins^22^.

Current methods of generating circular DNA are lengthy, inefficient, highly dependent on the length and sequence of DNA, and can result in unwanted chimeras^15,16^. The single-stranded DNA circularization is the commonest^17–24^. It is usually done using a splint (padlock probe) that is complementary to the ends of the DNA and hybridizes to it, bringing the two ends into close proximity in a double-stranded region for a ligase to join. This may also be done for small fragments without a splint using CircleLigase^29–31^. BGI’s nanoball technology uses splint circularization as follows: DNA is fragmented, and 100-300bp fragments are size selected; end repair, a cleanup, and the ligation of PCR adapters; a PCR to amplify the library using a phosphorylated primer; a second cleanup; and incubation of the products with a splint and a ligase (often, an exonuclease such as lambda would be used to degrade one DNA strand to limit rehybridization). The splint is then used as the primer for rolling circle amplification using phi29 polymerase – which has exquisitely high fidelity, is isothermal, and has vigorous strand displacement activity – to form “DNA nanoballs.” Here, we demonstrate that self-circularization of a large amplicon, and circularization using splints and T4 ligase, Taq DNA ligase or Ampligase, produce circular DNA but appear to have low efficiency. We also devised a method that utilizes asymmetric PCR to produce single-stranded DNA species without requiring exonuclease digestion of double-stranded DNA products (e.g., using lambda exonuclease). This methodology streamlines the process, eliminates several cleanups and a digestion, and reduces the time to result.

The incorporation of a double-stranded amplicon into a double-stranded DNA vector backbone is utilized in cloning. However, this approach has several drawbacks, including the presence of the vector backbone in all downstream applications. Here, we attempted to modify three common cloning techniques, Gibson and restriction enzyme cloning, and kinase ligase digestion, to self-circularize a double-stranded DNA amplicon. We could not generate singular, detectable circularized products using either approach. However, we produced several long concatemers, some of which may have been circular. This, however, is not desirable. We, too, were unable to produce singular, detectable circularized products by treating our amplicon as a vector backbone and ligating it to a short complementary fragment.

The production of pseudocircular DNA is least studied but is utilized by PacBio in their Single Molecule, Real-Time (SMRT) sequencing system. Here we successfully demonstrated a DumBell method, where single-stranded, phosphorylated hairpin DNA is ligated to blunt double-stranded phosphorylated PCR amplicons. While this method works well, circular DNA made only from DumBells and concatemers of amplicons are generated due to the blunt amplicons. A size selection DNA cleanup can remove circular DNA made only from DumBells while including unique molecular identifiers (short, random sequences) in the primers used for PCR, the concatemers can be bioinformatically deconvoluted^32^. Given this undesirable procedure and the artifacts generated, we developed two novel methods for the pseudo-circular of DNA and successfully demonstrated that they work as desired. First, uDumBell includes a deoxyUridine in the PCR primer sequences used to generate amplicons, which arrest elongation by Q5 and other high-fidelity polymerases before producing blunt DNA. The resulting 5’ overhangs allow for the ligation of complementary (non-blunt) dumbbell (hairpin) oligonucleotides. This ligation can be carried out in the same reaction buffer as the PCR, eliminating the need for DNA cleanups. This method can also be used to ligate any number of dsDNA fragments for cloning (**Supplementary Figure 2** - uQuickClone). Second, easyDB primers include a short tail sequence that forms a six-nucleotide hairpin at temperatures below 55°C, but not at temperatures above 55°C and thus, do not interfere with primer annealing. By including phosphorothioate bonds at the start of the complementary region in these primers, we inhibit T7 Exonuclease activity. Following PCR, the addition of T7 Exonuclease, a polymerase and a ligase, and incubation at 55°C or lower results in pseudo-circular double-stranded DNA (covalently closed circular DNA). We also made pseudo-circular DNA using TelN protelomerase. TelN was isolated from phage N15 and cuts dsDNA at a 56bp recognition sequence leaving covalently closed ends at the cleavage site. This method, however, requires the including a 56-bp tail on primer sequences, which interferes with PCR, and increases the cost of primer generation while decreasing its fidelity.

We demonstrate several methods, including using splints, a Gibson cloning-based approach for self-circularization, and novel methods for generating pseudo-circular DNA from a ~700bp amplicon of *rv0678*, the high GC content (65%) gene implicated in bedaquiline resistance in *M tuberculosis*, the causative agent of tuberculosis. This circular DNA can be used as a template for rolling circle amplification followed by long read sequencing, allowing for the error correction of sequence data^32^, and improving confidence in the resistance determination. The protection of circular DNA from degradation has applications in DNA vaccines, where DNA must be delivered into cells and make its way into the nucleus to assert its effects. Linear DNA with free ends is more recombinogenic^33^ and has lower transfection efficiencies and expression than DNA minicircles^34^ (dsDNA supercoiled circles containing only the genes of interest). The behavior of pseudo-circular DNA, however, is unknown. Pseudo-circular DNA is linear, double-stranded DNA with covalently closed (hairpin) ends and, unlike plasmids and minicircles, has no lower size limit. For these reasons, pseudo-circular DNA may have applications in transgenics or DNA vaccines.

## Acknowledgments

This study was funded by the National Institute of Allergy and Infectious Diseases (NIAID) 1R01AI153213-01A1. We have no conflicts of interest to declare.

## SUPPLEMENTARY MATERIAL

**Supplementary Figure 1:**
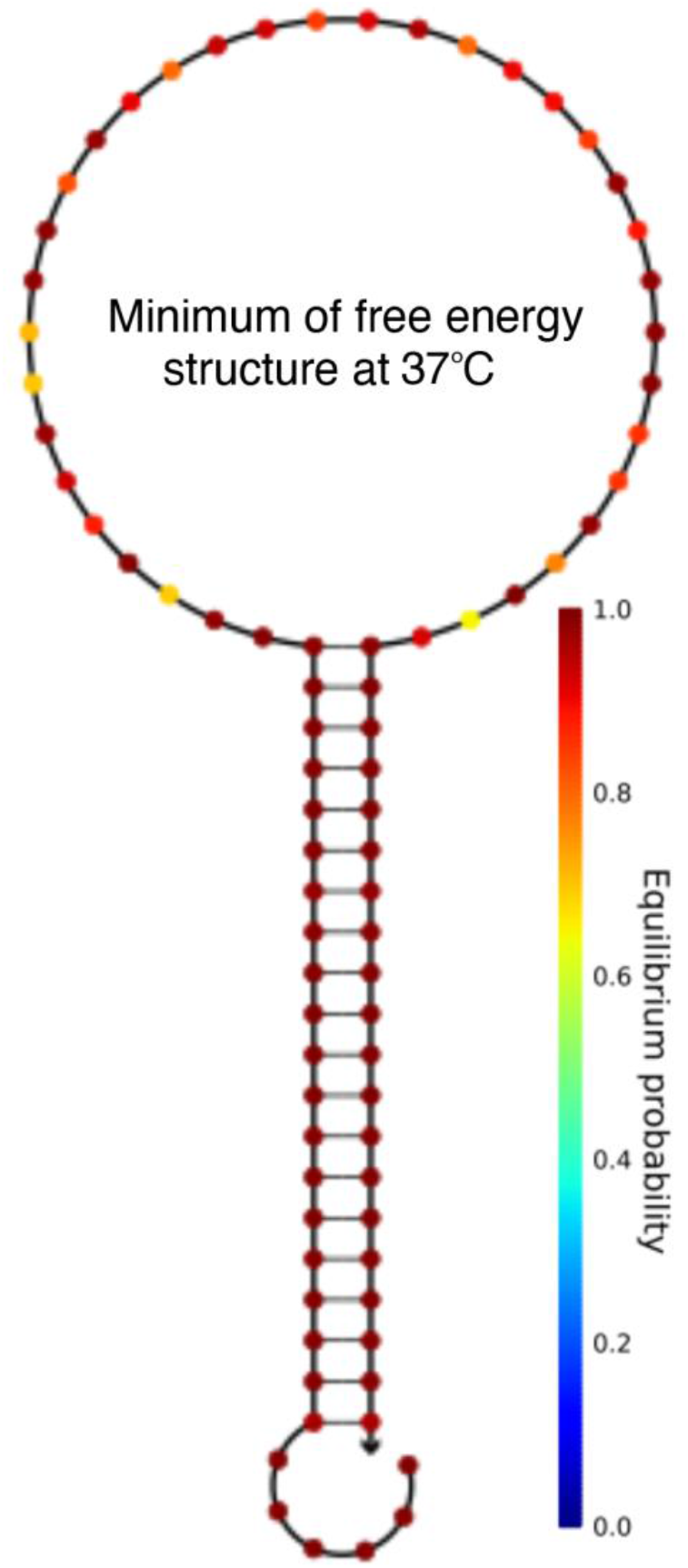
Structure of the uDumBell adapter sequence that forms a stable hairpin at temperatures ≤37°C.

**Supplementary Figure 2:**
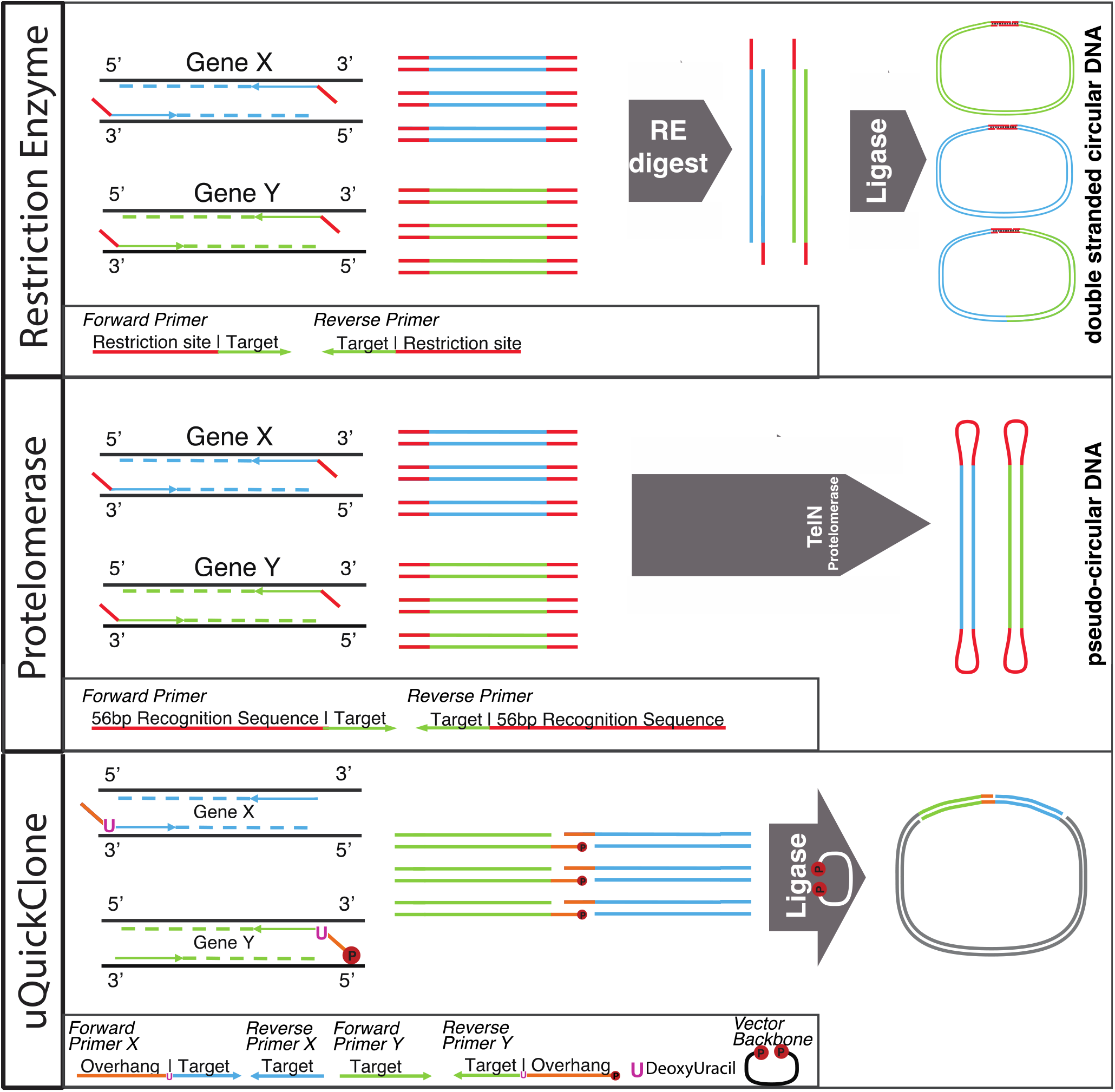
Streamlined methods for the circularization of PCR amplicons and the joining of amplicons with applications in cloning.

**Supplementary Figure 3:**
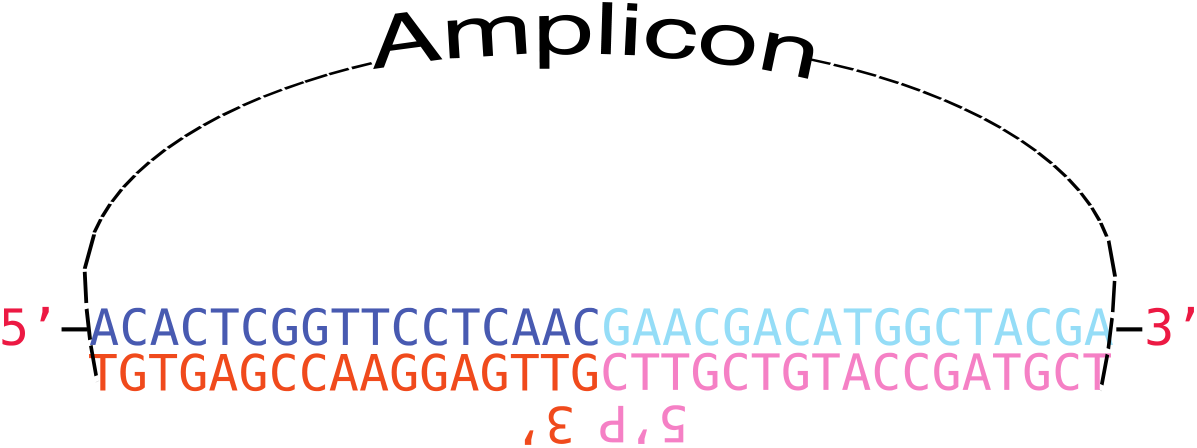
Structure of the single-stranded splint oligonucleotide (navy/blue) used in the splint method, bound to the single-stranded amplicon (red/pink).

**Supplementary Figure 4:**
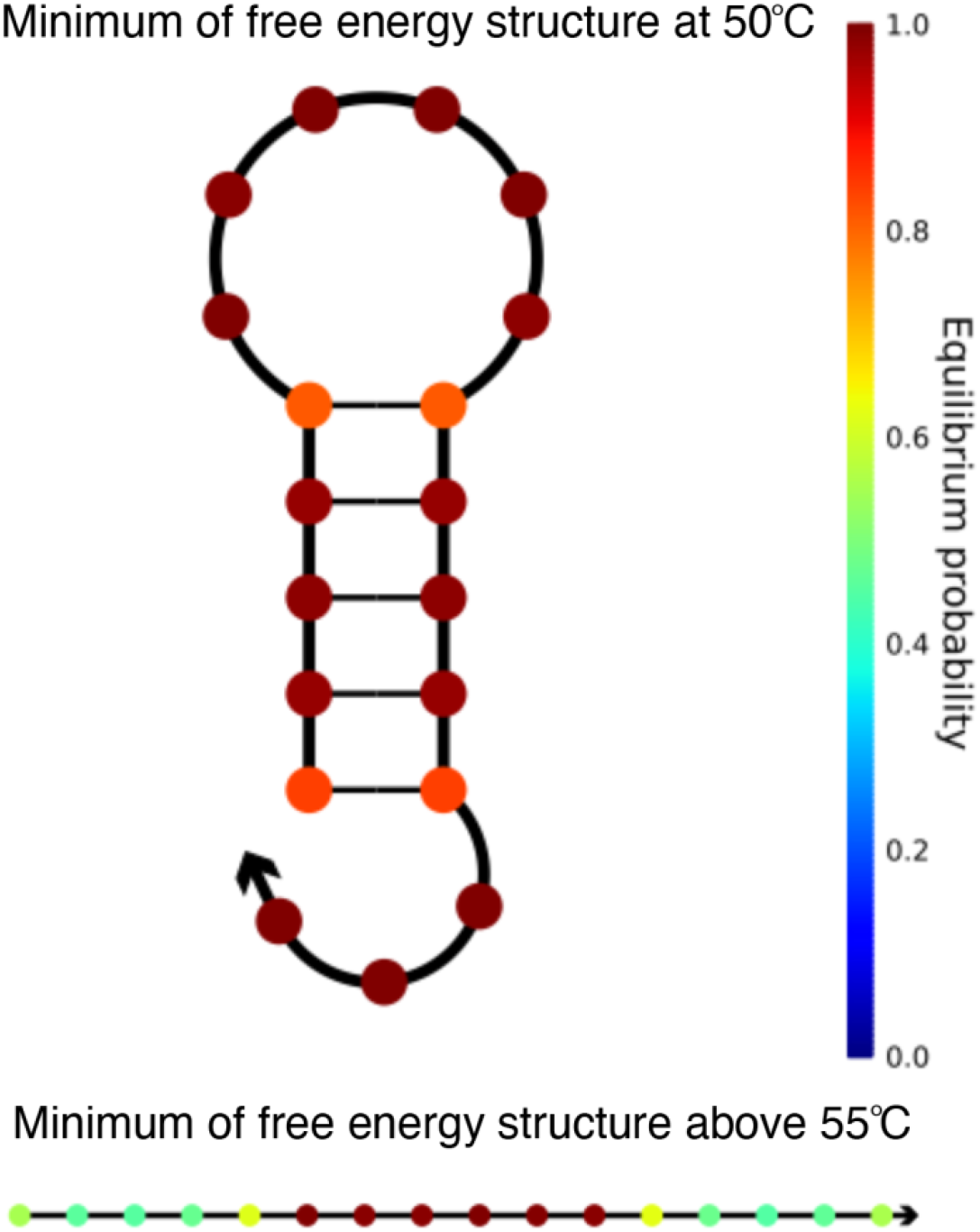
Structure of the tail sequence of the easyBD method that forms a hairpin at temperature <55°C, but not ≥55°C.

## References

1. Smith JH, Beals TP. Detection of nucleic acid targets using ramified rolling circle DNA amplification: a single nucleotide polymorphism assay model. PLoS One. 2013;8(5):e65053.

2. Konry T, Hayman RB, Walt DR. Microsphere-based rolling circle amplification microarray for the detection of DNA and proteins in a single assay. Anal Chem. 2009;81(14):5777–5782.

3. Larsson C, Grundberg I, Söderberg O, Nilsson M. In situ detection and genotyping of individual mRNA molecules. Nat Methods. 2010;7(5):395–397.

4. de la Torre TZG, Mezger A, Herthnek D, et al. Detection of rolling circle amplified DNA molecules using probe-tagged magnetic nanobeads in a portable AC susceptometer. Biosens Bioelectron. 2011;29(1):195–199.

5. Smolina I v, Cherny DI, Nietupski RM, et al. High-density fluorescently labeled rolling-circle amplicons for DNA diagnostics. Anal Biochem. 2005;347:152–155.

6. Østerberg FW, Rizzi G, Donolato M, et al. On-Chip Detection of Rolling Circle Amplified DNA Molecules from Bacillus Globigii Spores and Vibrio Cholerae. Small. 2014;10(14):2877–2882.

7. Rolling circle replication reporter systems. Published online January 2, 2002.

8. Daubendiek SL, Ryan K, Kool ET. Rolling-Circle RNA Synthesis: Circular Oligonucleotides as Efficient Substrates for T7 RNA Polymerase. J Am Chem Soc. 1995;117(29):7818–7819. doi:10.1021/JA00134A032/ASSET/JA00134A032.FP.PNG_V03

9. Methods for the isothermal amplification of nucleic acid molecules. Published online August 4, 1993.

10. Circular extension for generating multiple nucleic acid complements. Published online July 17, 1991.

11. Zhang DY, Brandwein M, Hsuih TC, H. NUCLEIC ACID AMPLIFICATION METHOD: RAMIFICATION-EXTENSION AMPLIFICATION METHOD (RAM). Published online February 5, 1998.

12. Volden R, Palmer T, Byrne A, et al. Improving nanopore read accuracy with the R2C2 method enables the sequencing of highly multiplexed full-length single-cell cDNA. Proc Natl Acad Sci U S A. 2018;115(39):9726–9731. doi:10.1073/PNAS.1806447115/SUPPL_FILE/PNAS.1806447115.SAPP.PDF

13. Pomerantz A, Peñafiel N, Arteaga A, et al. Real-time DNA barcoding in a rainforest using nanopore sequencing: opportunities for rapid biodiversity assessments and local capacity building. Gigascience. 2018;7(4):1–14. doi:10.1093/GIGASCIENCE/GIY033

14. Smith C, Halse TA, Shea J, et al. Assessing nanopore sequencing for clinical diagnostics: A comparison of Next-Generation Sequencing (NGS) methods for mycobacterium tuberculosis. J Clin Microbiol. 2021;59(1). doi:10.1128/JCM.00583-20/SUPPL_FILE/JCM.00583-20-S0007.XLSX

15. Ohyama T. MOLECULAR BIOLOGY INTELUGENCE UNIT DNA Conformation and Transcription. Accessed August 26, 2022. http://www.eurekah.com

16. de Santis P, Fuà M, Savino M, Anselmi C, Bocchinfuso G. Sequence dependent circularization of DNAs: A physical model to predict the DNA sequence dependent propensity to circularization and its changes in the presence of protein-induced bending. Journal of Physical Chemistry. 1996;100(23):9968–9976. doi:10.1021/JP9526096/ASSET/IMAGES/LARGE/JP9526096F00013.JPEG

17. Zhang DY, Brandwein M, Hsuih T, Li HB. Ramification amplification: a novel isothermal DNA amplification method. Mol Diagn. 2001;6(2):141–150. doi:10.1054/MODI.2001.25323

18. Xu H, Zhang Y, Zhang S, et al. Ultrasensitive assay based on a combined cascade amplification by nicking-mediated rolling circle amplification and symmetric strand-displacement amplification. Anal Chim Acta. 2019;1047:172–178. doi:10.1016/J.ACA.2018.10.004

19. Thomas DC, Nardone GA, Randall SK. Amplification of padlock probes for DNA diagnostics by cascade rolling circle amplification or the polymerase chain reaction. Arch Pathol Lab Med. 1999;123(12):1170–1176. doi:10.1043/1543-2165-123.20.1170

20. Tian B, Fock J, Minero GAS, Garbarino F, Hansen MF. Ultrasensitive Real-Time Rolling Circle Amplification Detection Enhanced by Nicking-Induced Tandem-Acting Polymerases. Anal Chem. 2019;91(15):10102–10109. doi:10.1021/ACS.ANALCHEM.9B02073/SUPPL_FILE/AC9B02073_SI_001.PDF

21. Wu L, Ling Y, Yang A, Wang S. Detection DNA point mutation with rolling-circle amplification chip. 2010 4th International Conference on Bioinformatics and Biomedical Engineering, iCBBE 2010. Published online 2010. doi:10.1109/ICBBE.2010.5518059

22. Schmidt TL, Beliveau BJ, Uca YO, et al. Scalable amplification of strand subsets from chip-synthesized oligonucleotide libraries. Nature Communications 2015 6:1. 2015;6(1):1–7. doi:10.1038/ncomms9634

23. Gong J, Li Y, Lin T, Feng X, Chu L. Multiplex real-time PCR assay combined with rolling circle amplification (MPRP) using universal primers for non-invasive detection of tumor-related mutations. RSC Adv. 2018;8(48):27375–27381. doi:10.1039/C8RA05259J

24. van Emmerik CL, Gachulincova I, Lobbia VR, et al. Ramified rolling circle amplification for synthesis of nucleosomal DNA sequences. Anal Biochem. 2020;588:113469. doi:10.1016/j.ab.2019.113469

25. Xu Y, Lin Z, Tang C, et al. A new massively parallel nanoball sequencing platform for whole exome research. BMC Bioinformatics. 2019;20(1):153. doi:10.1186/s12859-019-2751-3

26. Limberis J. uDumBell – Circularization of rv0678 for genotypic bedaquiline resistance testing of Mycobacterium tuberculosis. protocols.io. Published online 2022. doi:10.17504/protocols.io.36wgqj9zyvk5/v1

27. Jason Limberis. easyDB – Circularization of rv0678 for genotypic bedaquiline resistance testing of Mycobacterium tuberculosis. protocols.io. Published online 2022. doi:10.17504/protocols.io.3byl4j24rlo5/v1

28. Cameron CE, Götte M, Raney KD. Viral Genome Replication. Springer; 2009.

29. Polidoros AN, Pasentsis K, Tsaftaris AS. Rolling circle amplification-RACE: A method for simultaneous isolation of 5’ and 3’ cDNA ends from amplified cDNA templates. Biotechniques. 2006;41(1):35–42. doi:10.2144/000112205/ASSET/IMAGES/LARGE/FIGURE2.JPEG

30. Reiß E, Hölzel R, von Nickisch-Rosenegk M, Bier FF. Rolling Circle Amplification For Spatially Directed Synthesis Of A Solid Phase Anchored Single-Stranded DNA Molecule. AIP Conf Proc. 2006;859(1):25. doi:10.1063/1.2360583

31. Lou DI, McBee RM, Sawyer SL, et al. High-Throughput dna sequencing errors are reduced by orders of magnitude using circle sequencing. Proc Natl Acad Sci U S A. 2013;110(49):19872–19877. doi:10.1073/PNAS.1319590110/SUPPL_FILE/PNAS.201319590SI.PDF

32. Jason Limberis. RCAnalysis. github. Published online 2022. doi:10.5281/zenodo.7089302

33. Brinster RL, Chen HY, Trumbauer ME, Yagle MK, Palmiter RD. Factors affecting the efficiency of introducing foreign DNA into mice by microinjecting eggs. Proc Natl Acad Sci U S A. 1985;82(13):4438. doi:10.1073/PNAS.82.13.4438

34. Yasunobu K, Tomoyuki M, Haruji N, Kazuyuki K, Akihisa N, Fumio I. In vivo correlation between DNA supercoiling and transcription. Gene. 1981;13(2):173–184. doi:10.1016/0378-1119(81)90006-8

